# Herd immunity on chip: recapitulating virus transmission in human society

**DOI:** 10.1101/2022.05.27.493795

**Authors:** Wanyoung Lim, Narina Jung, Jiande Zhang, Zhenzhong Chen, Byung Mook Weon, Sungsu Park

**Affiliations:** Department of Biomedical Engineering, Sungkyunkwan University; Suwon, 16419, Korea; School of Advanced Materials Science and Engineering, Sungkyunkwan University; Suwon, 16419, Korea; School of Computational Sciences, Korea Institute for Advanced Study (KIAS); Seoul 02455, Korea; School of Mechanical Engineering, Sungkyunkwan University; Suwon, 16419, Korea; Research Center for Advanced Materials Technology, Sungkyunkwan University; Suwon, 16419, Korea; Institute of Quantum Biophysics (iQB), Sungkyunkwan University; Suwon, 16419, Korea

**Author notes:** Corresponding author. (B. M. Weon)/ (S. Park). These authors contributed equally to this work (Wanyoung Lim, Narina Jung).

## Abstract

Virus transmission is affected by population density, social distancing, and vaccination. This has been simulated only by mathematical models. Here, we report the first experimental model to mimic herd immunity to a human coronavirus using a microfluidic device filled with host cells. The device consists of 444 microchambers filled with susceptible (S_0_), infected (I_0_), and unsusceptible (U_0_) cells at specific ratios. The transmission rate and reproduction numbers were directly proportional to S_0_ and I_0_ and inversely proportional to U_0_. Herd immunity was achieved when the proportion of U_0_ was at 80% in a fixed number of uninfected (S_0_+U_0_) cells. These results were consistent with those from a mathematical model. The device can be used for predicting virus transmission.

**One-Sentence Summary:** We present the first experimental model enabling the simulation of herd immunity in a microfluidic device filled with host cells to human coronavirus.

## Main Text

Virus transmission is affected by the number of people in a city or country, contacts between people, social distancing, and the vaccination rate. The world is now entering the third year of the severe acute respiratory syndrome coronavirus 2 (SARS-CoV-2) pandemic, and each country has independently implemented preventative policies, such as wearing masks, social distancing, and lockdown, according to the number of confirmed cases and the vaccination rates. Due to the diversity of economic and social circumstances among populations, tools are needed for determining sustainable strategies to prevent SARS-CoV-2 transmission while avoiding as much disruption as possible. Mathematical models have been used to simulate various circumstances. The classical model of Susceptible-Infectious-Recovered (SIR) and its extensions have provided essential insights into the critical factors involved in controlling viral transmission dynamics through contact rates per individual (*1*). Furthermore, the social system can be simulated as a contact network with assumption-dependent variables for various situations, such as school, workplace, or age group. Based on the model-based predictions of the critical vaccination coverage, governments are working to increase vaccination rates to achieve herd immunity (*2, 3*). However, a model that can experimentally study virus transmission based on complex social behavior has not been established, and it is difficult to clearly define the parameter sets used for simulation of virus transmission.

A network of cells can be constructed in a microfluidic device by microfabrication technology (*4*) using the bacteria culture technique. We developed a virus transmission model on a microfluidic device consisting of 444 interconnected hexagonal microchambers (*5*) to create a so-called “herd immunity on chip (HIC)” to bridge the gap between simulations and biological conditions (**Fig. 1**). HIC consists of 444 interconnected hexagonal microchambers (diameter 200 μm, height 40 μm) with the largest gap of 30 µm, allowing cells to migrate into adjacent microchambers (**Fig. S1**). Its peripheral microchambers had sides of microslits with a gap of 5 µm, allowing nutrients to penetrate into the inner microchambers. An epicenter was punched (with a 700 μm diameter) to seed infected cells (I_0_) to infect initial susceptible cells (S_0_) (**Fig. 2A**). Thus, virus spread from the epicenter to the microchambers, and the number of microchambers containing infected cells increased daily. We used human coronavirus 229E (HCoV-229E) (*6*) to infect human lung fibroblast cells (MRC-5) as hosts as cytopathic effect such as rounding and aggregating of cells was observed by the virus (**Fig. S2**). To provide a quantitative comparison with the virus transmission on HIC, we extend the classical model of SIR (*1*) by considering five stages of cell infection; susceptible (S), expected (E), infected (I), and unsusceptible (U), as well as dead (D) cells, collectively termed SEIUD. The expected cells denote those that are infected but not yet infectious. The dynamics of cell infection in a device with multi-component chambers is different from that of the human infection in that a virus spreads through random motions of agents in a whole population. The virus transmission dynamics for an individual chamber on the chip is described by the chamber-index ‘*i*.’ Our SEIUD model includes cell divisions of susceptible and immune cells but does not consider recovered cells because cells typically die within one day after being infected, which is shorter than the span of virus transmission. We also assume that there is no cell division for the infected cells in our experiments.

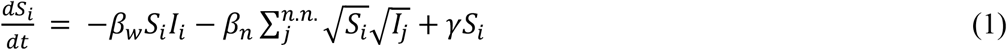

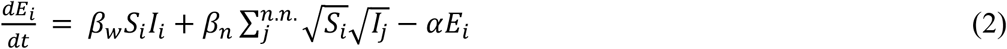

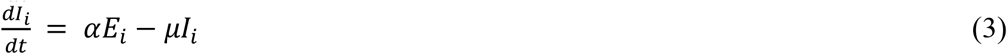

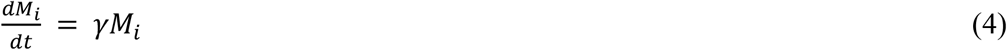

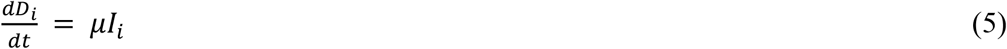

Here, *γ*is the division rate, *α* is the transition rate from expected to infected, and µ is the death rate. The model distinguishes the number of contacts of cells within each chamber from those across chambers using different contact rates: *β*_*w*_ is the average number of contacts per cell per time within a chamber, and *β*_n_ is that between nearest neighbor chambers. It is likely that the contacts of susceptible cells can be reduced in the presence of immune cells in a confined domain. We assume that the contact rates decrease with the population of immune cells when its population becomes larger than a threshold value *M*_*cr*_ : *β*_*w*_ = *β*_0_(1 − *c*(*M*_*i*_ *− M*_*cr*_)^*m*^) and *βn* = *β*_1_(1 − *c*(*M*_*i*_ *− M*_*cr*_)^*m*^). In the classical SIR model (*1*), it is assumed that the individual contact rates are determined based on excellent mixing among individuals. However, the contact rates of cells between nearest chambers are strictly constrained by the division of chambers in which cells in two different chambers will not be mixed with each other over time. Considering this limited cell motility between chambers, we assumed that the contacts mainly occurred at the interfacial regions. The corresponding cells contacted at the interfaces were counted with consideration of the square roots in the model. Parameters in the model, including the transition rates and division rate of cells, were determined by the gradient descent algorithm to minimize their fitting errors.

**Fig. 1.**
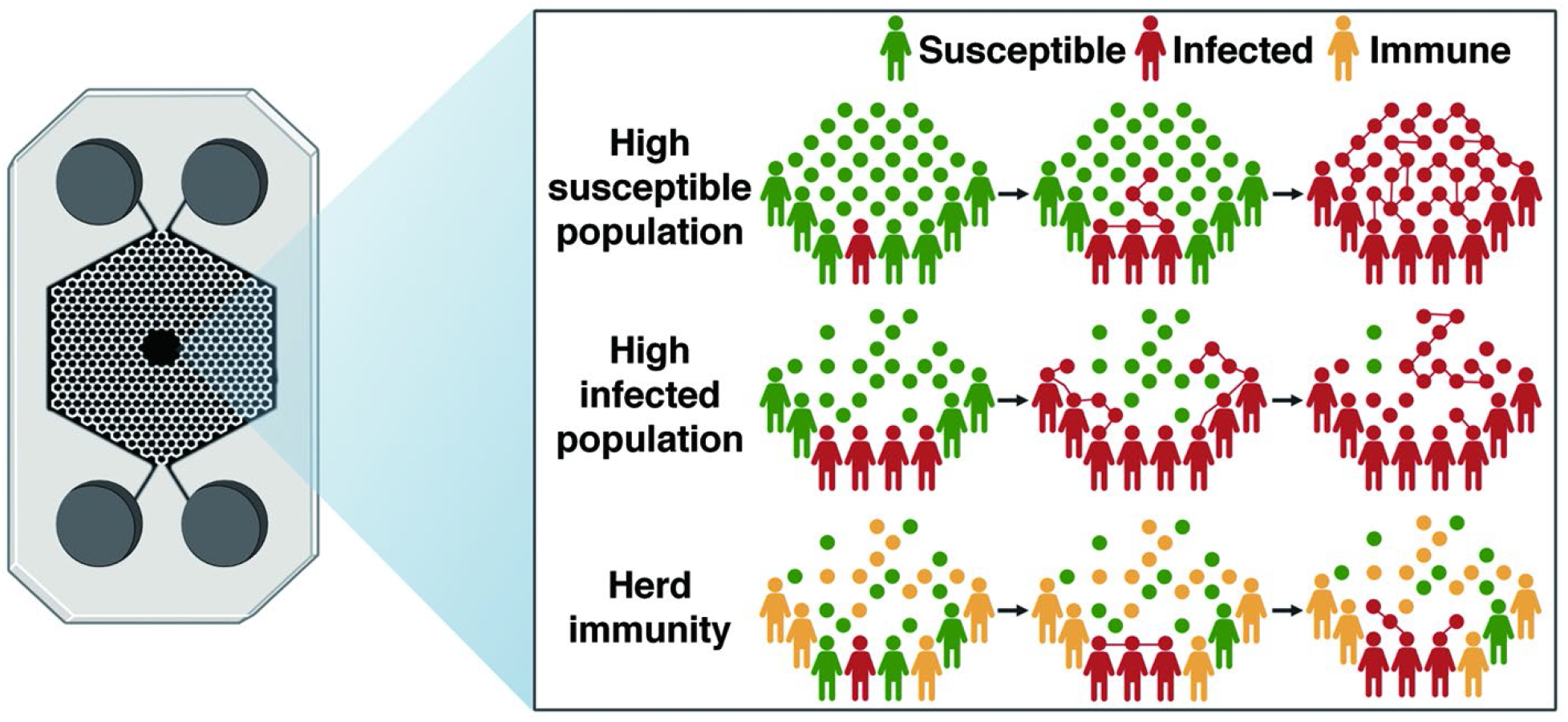
Schematic of recapitulating virus transmission in human society using herd immunity on chip (HIC). Human cells in green, red, and yellow are susceptible (S_0_), infected (I_0_), and unsusceptible (U_0_), respectively.

**Fig. 2.**
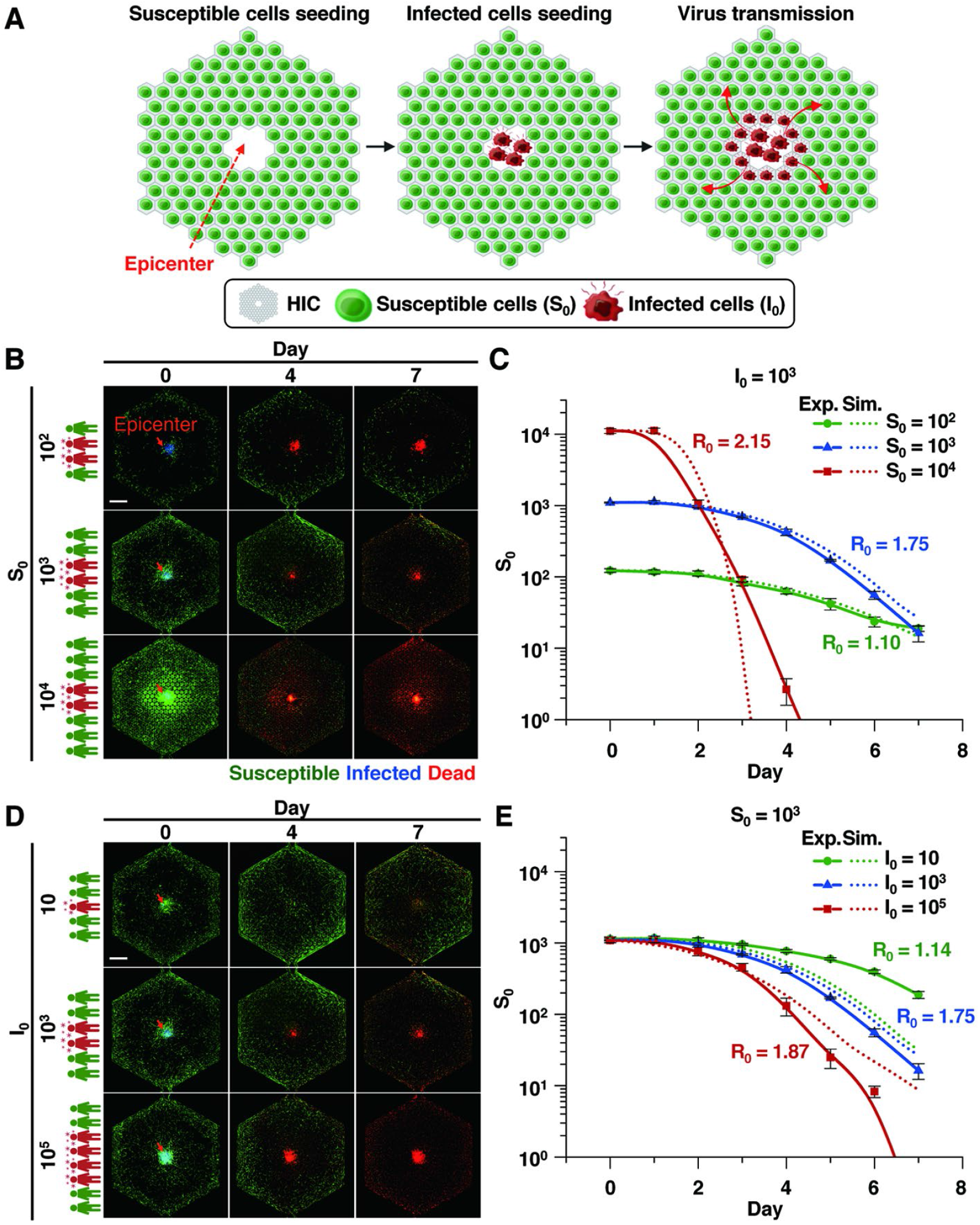
Virus transmission on HIC at different densities of susceptible (S_0_) and infected (I_0_) cells for 7 days. **(A)** Schematic of a process of virus transmission on HIC. **(B)** Different numbers (10^2^, 10^3^ and 1 0^4^) of S_0_ cells at I_0_ cells= 10^3^. **(C)** The S_0_ number (log) according to S_0_. **(D)** Different numbers (10^2^, 10^3^ and 1 0^4^) of S_0_ at I_0_ = 10^3^. Effect of I_0_ (10, 10^3^ and 10^5^) at S_0_ = 10^3^ on virus transmission. **(E)** The S_0_ number (log) for 7 days according to I_0_. Scale bar = 1 mm. Exp. = experimental results. Sim. = simulation results.

The transmission rate and reproduction number (R_0_) at I_0_ = 10^3^ increased as S_0_ increased from 10^2^ to 10^4^ (**Fig. 2B**). The average distance between cells in the microchambers of HIC was largest at S_0_ = 10^2^. Therefore, the number of contacts between cells was lowered, as was the probability of virus propagation. In contrast, at S_0_ = 10^4^, the cells were densely packed in microchambers, which allowed susceptible cells to be infected, and S_0_ was greatly reduced on day 4 (**Fig. 2C**). The R_0_ also decreased from 2.15 to 1.10 as S_0_ decreased. The reproduction numbers, which were all larger than one, increased as the population densities increased since contact rates of infected cells are proportional to population density (*7*). The experimental results using HIC were consistent with those from our SEIUD model. The transition rates of cells used in the computation were *β*_0_ = 0.132, *β*_1_ = 0.077, and *α* = 9.763 ; the division rate was *γ*= 0.048; the dealth rate was µ= 0.211; the parameters related to the immune response were *M*_*cr*_ = 2.105, *m* = 0.348, and *c* = 1.010. Our findings are also consistent with previous reports that have demonstrated a strong relationship between population density and other infectious diseases (*8-11*). Sy *et al*. (*12*) determined the association between population density and R_0_ of SARS-CoV-2 across the U.S. Using a densely populated city like New York City as an example set, the previous findings suggest that a greater population density facilitates more frequent interactions between susceptible and infectious individuals, sustaining the continued transmission and spread of COVID-19.

A similar trend was observed when I_0_ increased from 10 to 10^5^ at S_0_ = 10^3^ (**Fig. 2D**). As I_0_ increased, the intensity of blue fluorescence, which indicates I_0_, increased (**Fig. S3**). In the beginning, the total number of contacts between I_0_ and S_0_ was low at I_0_ = 10 cells, so the infection spread slowly. However, the total contacts between I_0_ and S_0_ were very high at I_0_ = 10^5^ cells, and S_0_ dramatically decreased. The delay in virus transmissions as we reduced I_0_ is demonstrated by the R_0_ decreased from 1.87 to 1.14 with I_0_ decrease (**Fig. 2E**). The experimental results were in line with the results of the SEIUD model. When I_0_ = 10, the minimum infective dose was low due to the structure of chambers and did not seem to be reflected in the simulation. However, the infected virus eventually spread throughout the chip. On February 9, 2020, 5,006 COVID-19 cases were confirmed in association with a church cluster in Daegu, South Korea. Prior to that date, South Korea had a low number of confirmed cases and appeared to have the outbreak under control (*13*). However, the transmission rate of COVID-19 increased by approximately 20-fold, and COVID-19 rapidly spread nationwide. The outbreaks highlight the potential for a high transmission rate of diseases with a larger number of infected cases. Taken together, our results suggest that controlling the number of infected people in the early stages can slow the spread of the virus.

To mimic herd immunity (**Fig. 3A**), HIC was first filled with different proportions (60– 80%) of unsusceptible cells (U_0_) in a fixed number (10^3^) of uninfected (S_0_ + U_0_) cells and later inoculated with 10^3^ of I_0_ in the epicenter (**Fig. 3B**). Aminopeptidase N (APN) knockdown MRC-5 cells, in which the APN receptor for virus binding is downregulated, were used as U_0_ (**Fig. S4**). Most S_0_ cells were infected within 7 days when the proportions of U_0_ cells were 60% and 70%. However, herd immunity was achieved when the proportion of U_0_ cells was 80% (**Fig. 3B, C**). Infection only occurred in the epicenter due to the close contact between infected and susceptible cells; the number of contacts dropped, and new infection events were prevented (**Fig. 3B**). The transmission rate and R_0_ decreased as U_0_ increased. The experimental results matched the results of the SEIUD model (**Fig. 3C**). Herd immunity could not be achieved when U_0_ was 80% in 96-well plates due to the lack of controllability in transmission (**Fig. S5**). Similar trends of virus transmission were also observed when other susceptible cells (Huh-7; a hepatocarcinoma cell line) (**Fig. S6**) and virus strain (human coronavirus OC43) (**Fig. S7**). In the COVID-19 pandemic, people have been uncertain about the success of the vaccines. Researchers worldwide are exploring many aspects, policies, and temporal predictions of this disease (*14-16*), all of which require effective mathematical modeling and simulations. Mondal *et al*. (*3*) showed that 50–60% of vulnerable patients might die before attaining approximately 70% immunization of the total population. Britton *et al*. (*17*) estimated that if R_0_ = 2.5 in an age-structured community with mixing rates fitted to social activity, the disease-induced herd immunity level would reach approximately 43%. However, predictions using simulations of the herd immunity threshold, that is, the minimum fraction of the population that needs to be immunized to eradicate the disease, vary due to heterogeneity of countries, cities, and populations. Recent studies reported that immunity levels should consider heterogeneities in continuously varying susceptibilities (*18, 19*). Our experimental model reflects the effect of such changes in HIC.

**Fig. 3.**
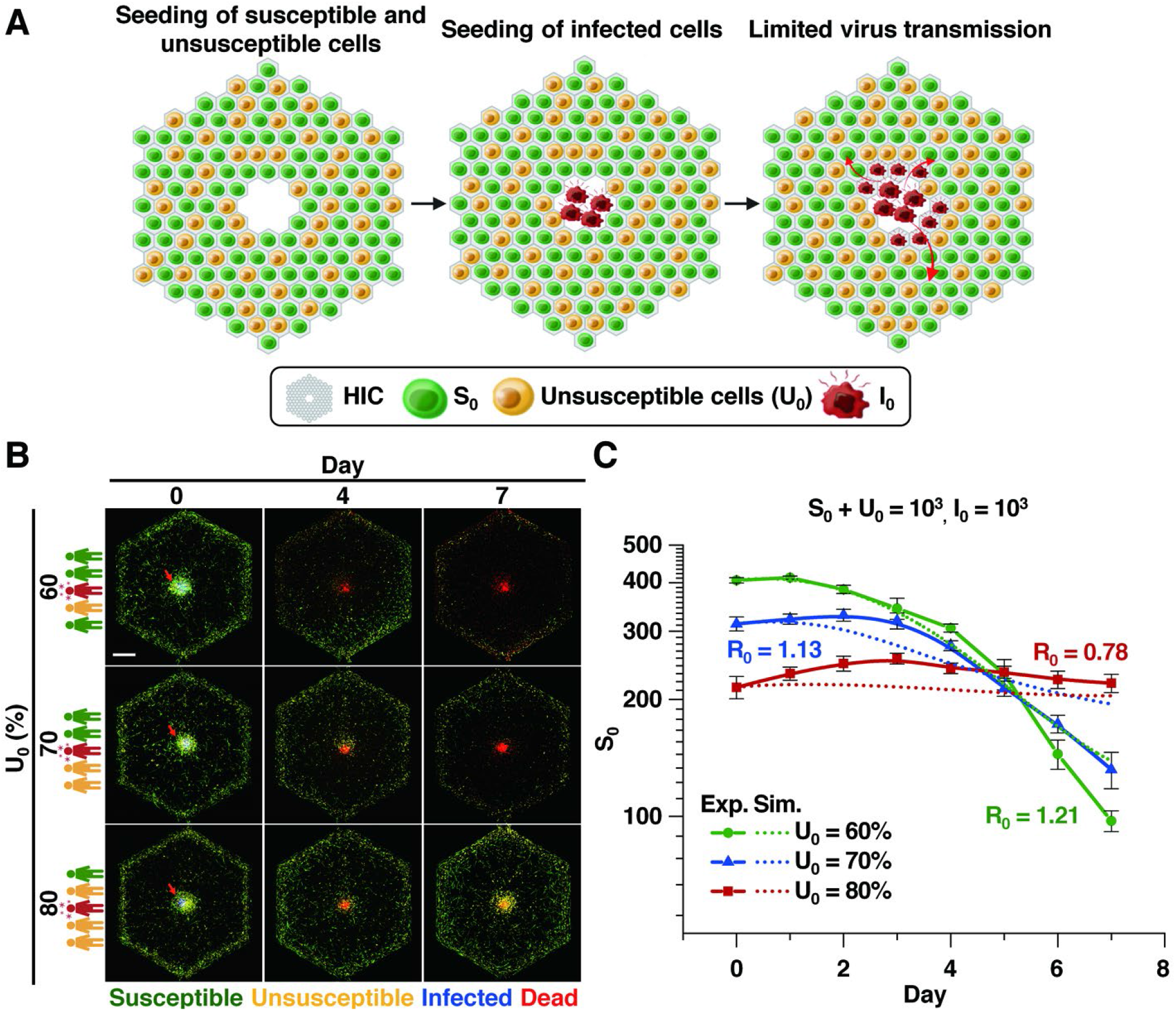
Effect of unsusceptible (U_0_) populations on virus transmission. **(A)** Schematic of a process of limited virus transmission on HIC. **(B)** Effect of the proportion (60%, 70%, and 80%) of U_0_ at S _0_ + U _0_ = 1 0^3^ a nd I_0_ = 10^3^ on virus transmission. **(C)** The S_0_ number (log) for 7 days according to U_0_. Scale bar = 1 mm.

We present the first experimental model enabling the simulation of herd immunity in a microfluidic device filled with host cells susceptible to human coronavirus. Our study demonstrates that HIC has the potential to be a powerful tool for predicting the transmission of viral diseases.

## Funding

Technology Innovation Program (20008413) funded by the Ministry of Trade, Industry & Energy (MOTIE)

BioNano Health-Guard Research Centre as a Global Frontier Project (H-guard 2018M3A6B2057299) through the National Research Foundation (NRF) of Ministry of Science and ICT (MSIT)

Basic Science Research Program and through the National Research Foundation of Korea (NRF) funded by the Ministry of Education (2016R1D1A1B01007133)

## Author contributions

Conceptualization: SP, WL

Experiments: WL, NJ, JZ, ZC

Funding acquisition: SP, BMW, NJ

Supervision: SP, BMW

Writing: SP, WL, BMW, NJ

## Competing interests

Authors declare that they have no competing interests.

## Data and materials availability

All data are available in the main text or the supplementary materials.

## Supplementary Materials

Materials and Methods

Figs. S1 to S7

References (*20–26*)

## Supplementary Materials for

## Materials and Methods

### Cell culture and viruses

The human fetal lung fibroblast cell line MRC-5 and human hepatocarcinoma cell line Huh-7 were purchased from Korean Cell Line Bank (Seoul, Korea), respectively. MRC-5 cells labeled with green fluorescent protein (MRC-5 GFP) were purchased from Lugen Sci Co., Ltd. (Bucheon, Korea). MRC-5 cells were maintained in minimum essential media (MEM) (HyClone Laboratories, Logan, UT) supplemented with 10% (v/v) fetal bovine serum (FBS) (HyClone Laboratories), 100 units penicillin mL^-1^ (Life Technologies, Carlsbad, CA), 100 μg streptomycin mL^-1^ (Life Technologies), 25 mM sodium bicarbonate (Sigma-Aldrich), and 25 mM hydroxyethyl piperazine ethane sulfonic acid (HEPES) (Life Technologies) at 37 °C under 5% CO_2_. Huh-7 cells were maintained in RPMI (HyClone Laboratories) supplemented with 10% (v/v) FBS, 100 units penicillin mL^-1^, and 100 μg streptomycin mL^-1^.

Stocks of human coronaviruses, human coronavirus 229E (HCoV-229E) and HCoV-OC43, were kindly provided by the Korea Research Institute of Bioscience and Biotechnology (KRIBB) (Daejeon, Korea) and Korea Bank for Pathogenic Viruses (KBPV) (Seoul, Korea), respectively.

### Determination of 50% tissue culture infective dose (TCID_50_)

TCID_50_ was used to determine virus infectivity (*20, 21*). In brief, freshly cultured MRC-5 or Hu7 cells (10^4^ cells per well) were seeded in a 96-well plate (Corning) and incubated at 37 °C under 5% CO_2_ overnight. Cells were first washed three times with phosphate-buffered saline (PBS) (pH 7.4) and then inoculated with HCoV-229E at different concentrations (10^−3^-10^3^ TCID_50_/mL) in MEM. The plates were incubated at 33 °C under 5% CO_2_ until cytopathic effect (CPE) such as rounding and aggregating of cells was observed. TCID_50_ of HCoV-OC43 was determined using the same method.

### Fabrication of HIC

The design of our previous chip (*5*) was slightly modified for HIC (**Fig. S1**). A silicon mold of HIC was fabricated using the negative photoresist SU–8 (MicroChem Corp., Westborough, MA) by photolithography (*22*). A replica layer of polydimethylsiloxane (PDMS) (Sylgard® 184, Dow Silicones Corp.) was then fabricated in the mold using soft lithography (*22, 23*). In details, a mixture of the PDMS prepolymer and curing agent in a 10:1 (w:w) ratio was cast onto the mold and cured at 80 °C for 2 h. Then, the cured PDMS layer was peeled off the mold. Afterward, an epicenter (diameter 0.7 mm) and four reservoirs (diameter 8 mm) were punched through the PDMS layer. The punched layer was finally bound to a cover glass after being treated with O_2_ plasma for 30 s. Dimensions of the completed HIC are 20 mm (width) × 30 mm (length) × 5 mm (height) (**Fig. S1**).

### Preparation of S_0_, I_0_ and U_0_ for cell seeding into HIC

MRC-5 labeled with GFP or Huh-7 labeled with the green fluorescent dye PKH67 (Sigma-Aldrich) was used as S_0_ for easy monitoring of cell seeding and virus transmission in HIC. In detail, approximately 5 × 10^5^ S_0_ were seeded in a cell culture flask and incubated at 37 °C under 5% CO_2_ until they formed a monolayer. Cells were then detached from the flask by trypsinization. Detached cells were suspended in culture medium right before cell seeding into HIC.

MRC-5 or Huh-7 was used to obtain I_0_. Before inoculation, cells in a monolayer were washed with PBS and inoculated with HCoV-229E or HCoV-OC43 at 100 TCID_50_/mL (final conc.) (*20*,). Afterward, they were incubated at 33 °C under 5% CO_2_ overnight (*21, 24*). Next day, the nuclei of I_0_ were labeled with Hoechst 33342 (Sigma-Aldrich).

To generate unsusceptible cells (U_0_), MRC-5 or Huh-7 was transfected with either aminopeptidase (APN) small interfering RNA (siRNA) oligonucleotide (sense: 5’-GCA GCA GAU CUG UAU AUU U-3’, antisense: 5’-AAA UAU ACA GAU CUG C-3’) or negative control siRNA from Bioneer Co. (Daejeon, Korea). The transfection efficiency was determined using quantitative real-time PCR (qRT-PCR). The primer sequences used were as follows: APN: sense, 5’-GTT CTC CTT CTC CAA CCT CAT C-3’, antisense, 5’-CTG TTT CCT CGT TGT CCT TCT-3’; glyceraldehyde-3-phosphate dehydrogenase (GAPDH): sense, 5’-AGG TGG TGA AGC AGG CGT CGG AGG G-3’, antisense, 5’-CAA AGT GGT CGT TGA GGG-3’ (*25*). Reduced virus infection in APN-knockdown (APN-KD) cells were confirmed before cell loading into HIC. Cells were labeled with CellMask™ Deep Red Plasma Membrane Stain (Invitrogen, Waltham, MA) prior to being loaded into HIC.

### Loading of S_0_, I_0_ and U_0_ into HIC and a 96-well plate

To study the effect of S_0_ density on virus transmission, different numbers (10^2^-10^4^) and a fixed number (10^3^) of S_0_ and I_0_ were loaded into HIC at different days. In detail, the hexagonal microchambers in HIC was loaded with S_0_ through the epicenter in the center of the chip using a pipette. The chip was then incubated at 37 °C under 5% CO_2_ for 1 day. The next day, about 10^3^ I_0_ were gently loaded into the epicenter by gravity-driven flow from a 300 µL tip containing them. On the other hand, to study the effect of I_0_ density on virus transmission, a fixed number (10^3^) and different numbers (10, 10^3^ and10^5^) of S_0_ and I_0_ were loaded into HIC in a similar manner. On the other hand, a total of 10^3^ cells composed of S_0_ and U_0_ in different proportions (4:6, 3:7 and 2:8) and 10^3^ I_0_ were loaded into HIC at different days to study the effect of U_0_ ratio on virus transmission. Once the cell loading was completed, the four reservoirs in HIC were filled with 200 μL of medium to provide cells in HIC with nutrients (*5*). The chip was then incubated at 33 °C under 5% CO_2_ for 7 days. Each day, 200 μL of fresh medium containing 4 μM ethidium homodimer-1 (EthD-1) was exchanged to stain dead cells by infection. To generate cell number data, optical and fluorescence images were obtained daily using a K1-Fluo confocal microscope (Nanoscope Systems, Inc., Daejeon, Korea). We employed the K1-array viewer to acquire stitched images across various channels at 10× magnification. Images were processed using Image J (NIH, Bethesda, MD). The number of live S_0_ was manually counted from day 0 to day 7.

For control experiments on HIC, S_0_, I_0_ and U_0_ were similarly seeded into a 96-well plate.

### Calculation of reproduction number (R_0_)

The reproduction number R_0_ for an SIR model is calculated as the product of transmission rate and infectious period (*26*). Here, the transmission rate was defined as an average of the daily increase rates of the infected chamber, and the infectious period was the time to show CPE in the chambers.

### Statistical analysis

Statistical significance was determined using a Student’s *t*-test. The results are presented as the mean ± standard deviation (S.D.). All quoted *p* values were two-tailed and differences were considered statistically significant at ∗*p* < 0.05: ∗∗*p* < 0.01; ∗∗* *p* < 0.001.

**Fig. S1.**
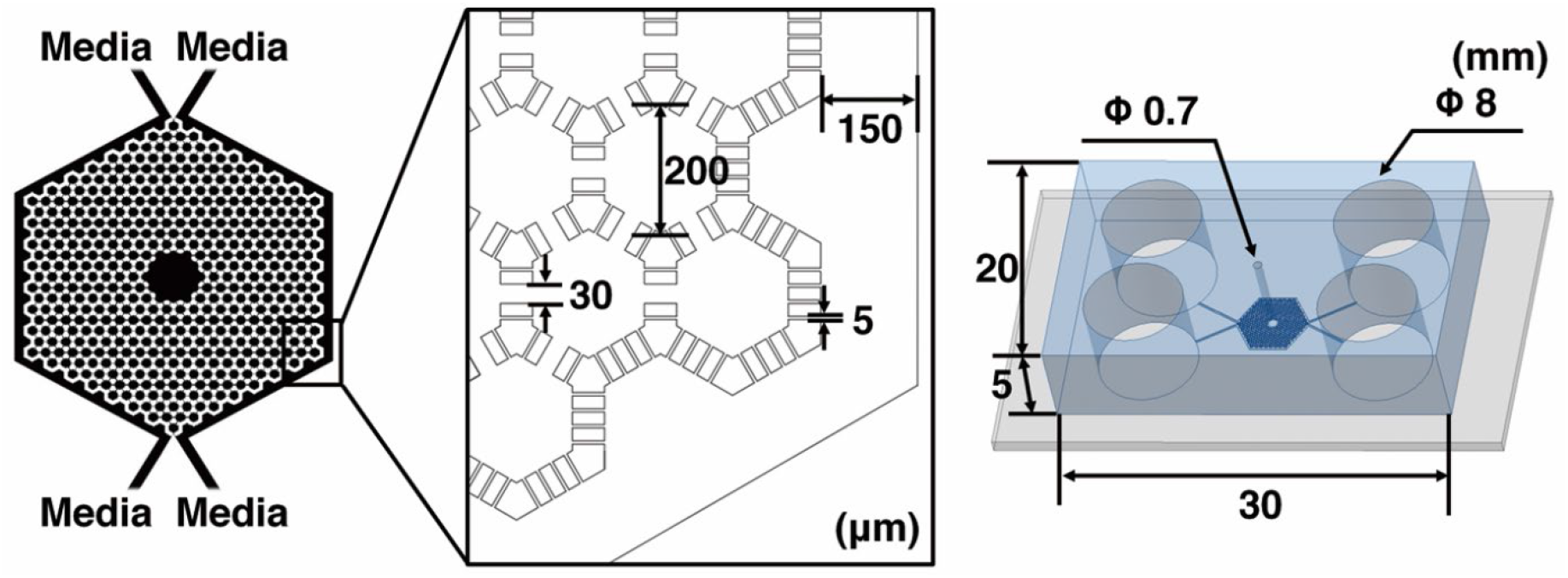
Schematic figures of HIC showing dimensions. HIC consists of 444 interconnected hexagonal microchambers (diameter 200 μm, height 40 μm) with the largest gap of 30 µm. An epicenter (diameter 0.7 mm) was fabricated to load cells, while four reservoirs (diameter 0.8 mm) were fabricated to provide cell culture media to the microchambers by gravity-driven flow through two microchannels surrounding the microchambers.

**Fig. S2.**
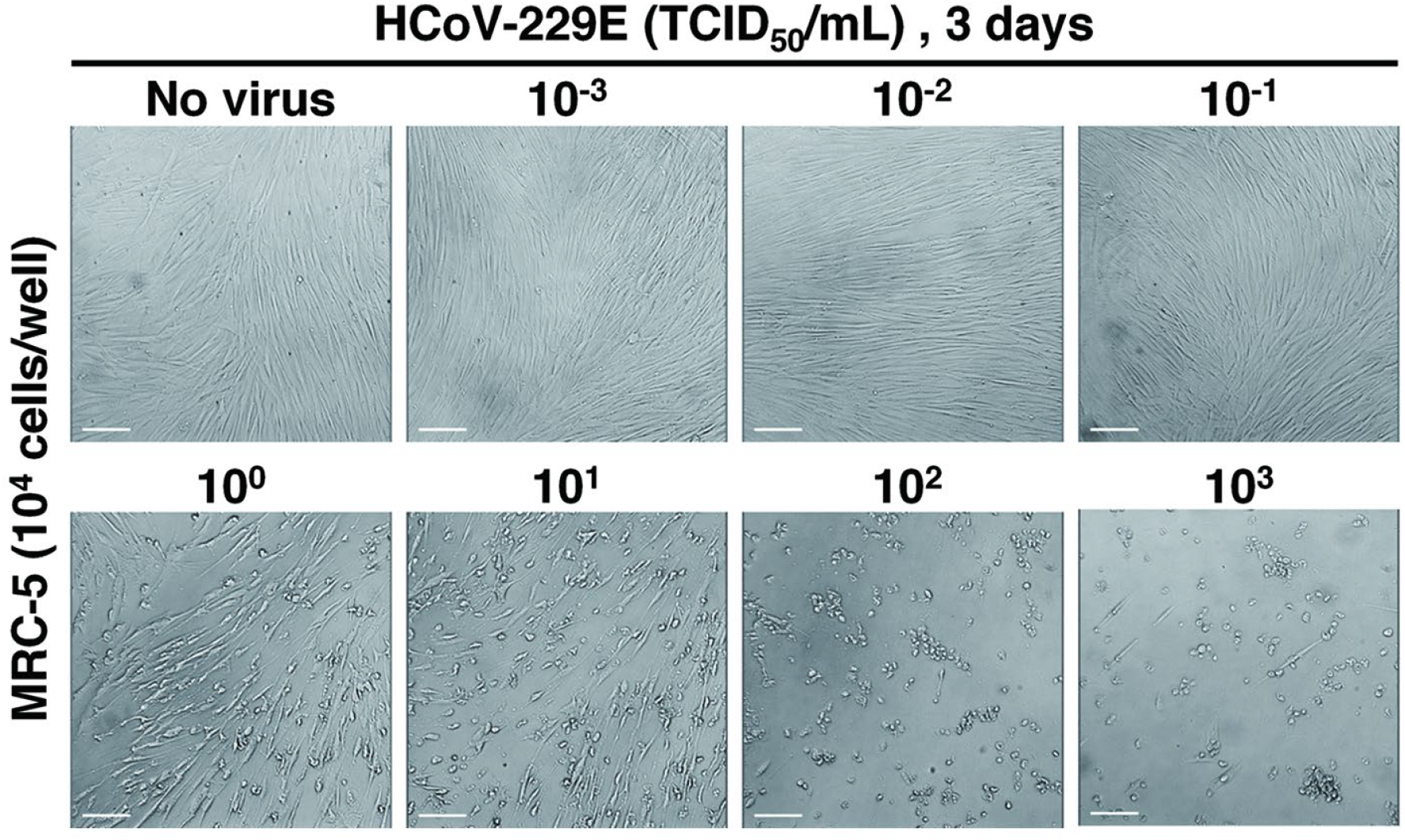
Morphological changes of MRC-5 cells after HCoV-229E infection. Cells were inoculated with 100 μL of HCoV-229E at different concentrations (10^−3^ to 10^3^ TCID_50_/mL) for 3 days. Cytopathic effect was used to determine TCID_50_. Scale bars = 100 μm.

**Fig. S3.**
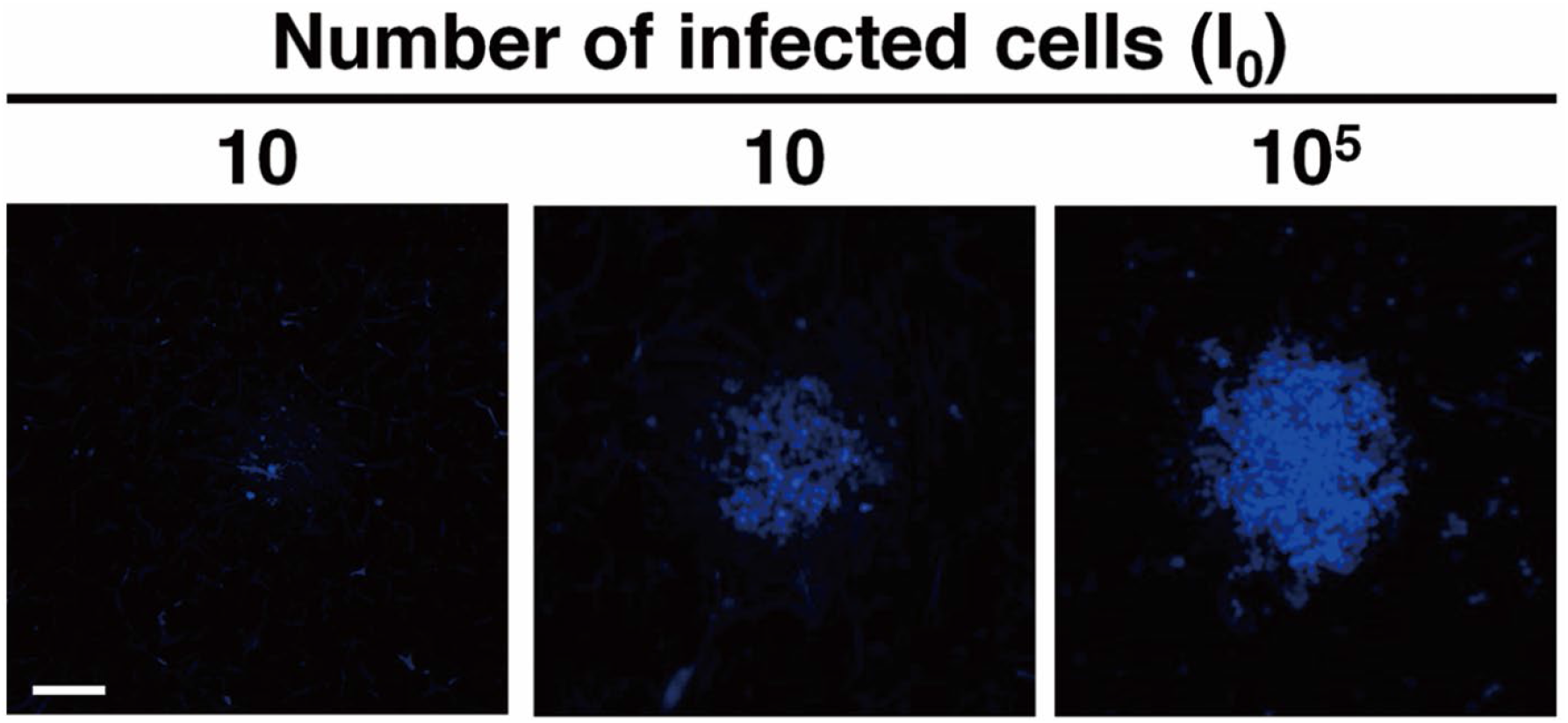
Fluorescent images of the epicenter (diameter 0.7 mm) in HIC after seeding of infected cells. Various numbers of infected cells (10, 10^3^ and 10^5^) were stained with Hoechst 33342 (blue) for 5 min and seeded into the epicenter. The images showed that most of inoculated cells remained in the epicenter. Scale bar = 200 μm.

**Fig. S4.**
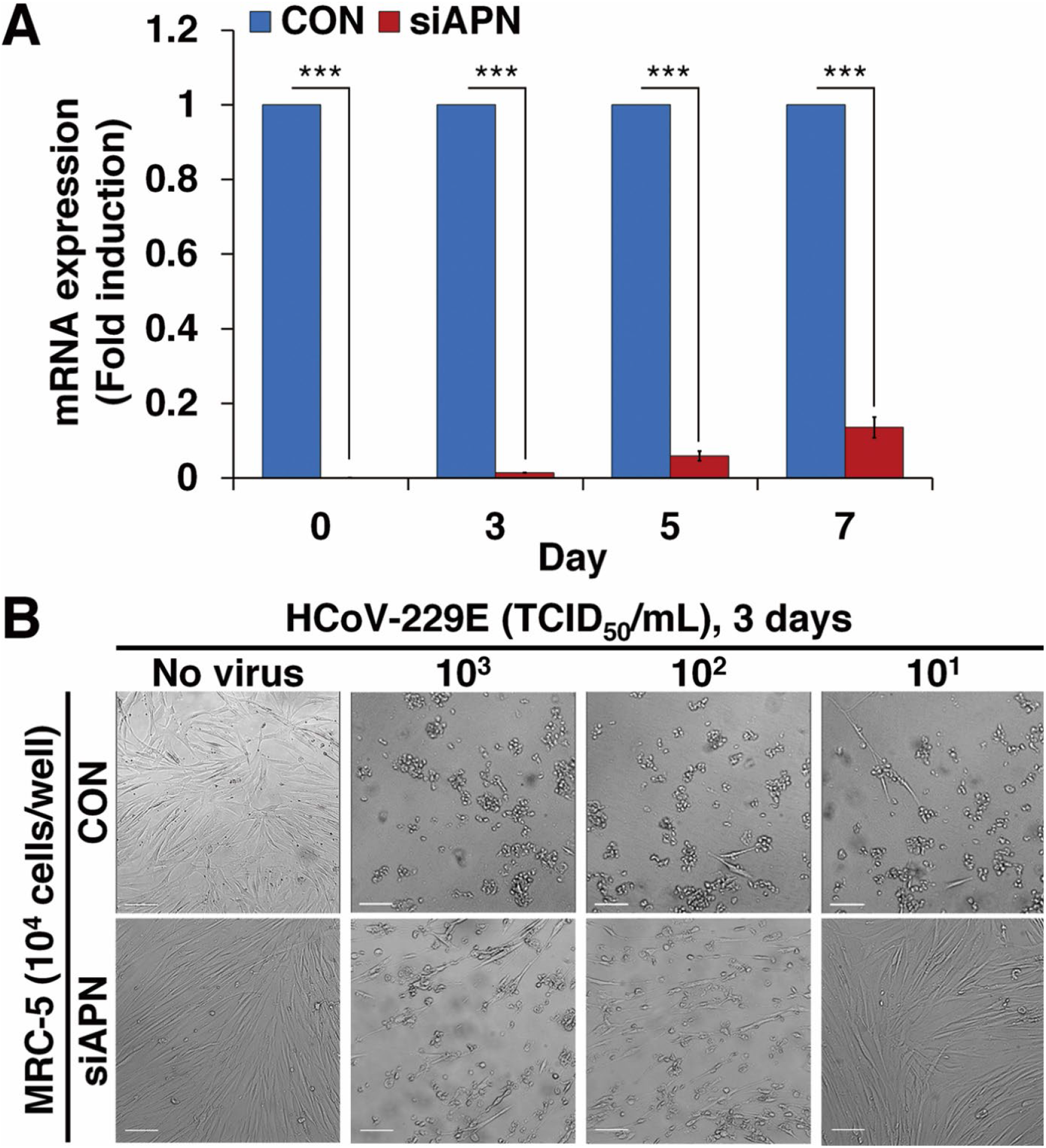
siRNA mediated inhibition of APN expression and virus infection in MRC-5 cells. **(A)** qRT-PCR analysis of APN mRNA expression at different days (0, 3, 5 and 7). CON: Control cells treated with negative control siRNA. siAPN: cells treated with siAPN. All the experiments were performed in triplicate. Data represent the mean ± S.D. Student’s *t*-test, \*\*\**p* < 0.001. **(B)** Morphological changes of CON and siAPN cells after HCoV-229E infection. Both types of cells were first inoculated with 100 μL of HCoV-229E at different concentrations (10^1^-10^3^ TCID_50_/mL) and then incubated at 33°C under 5% CO_2_ for 3 days. Scale bar = 100 μm.

**Fig. S5.**
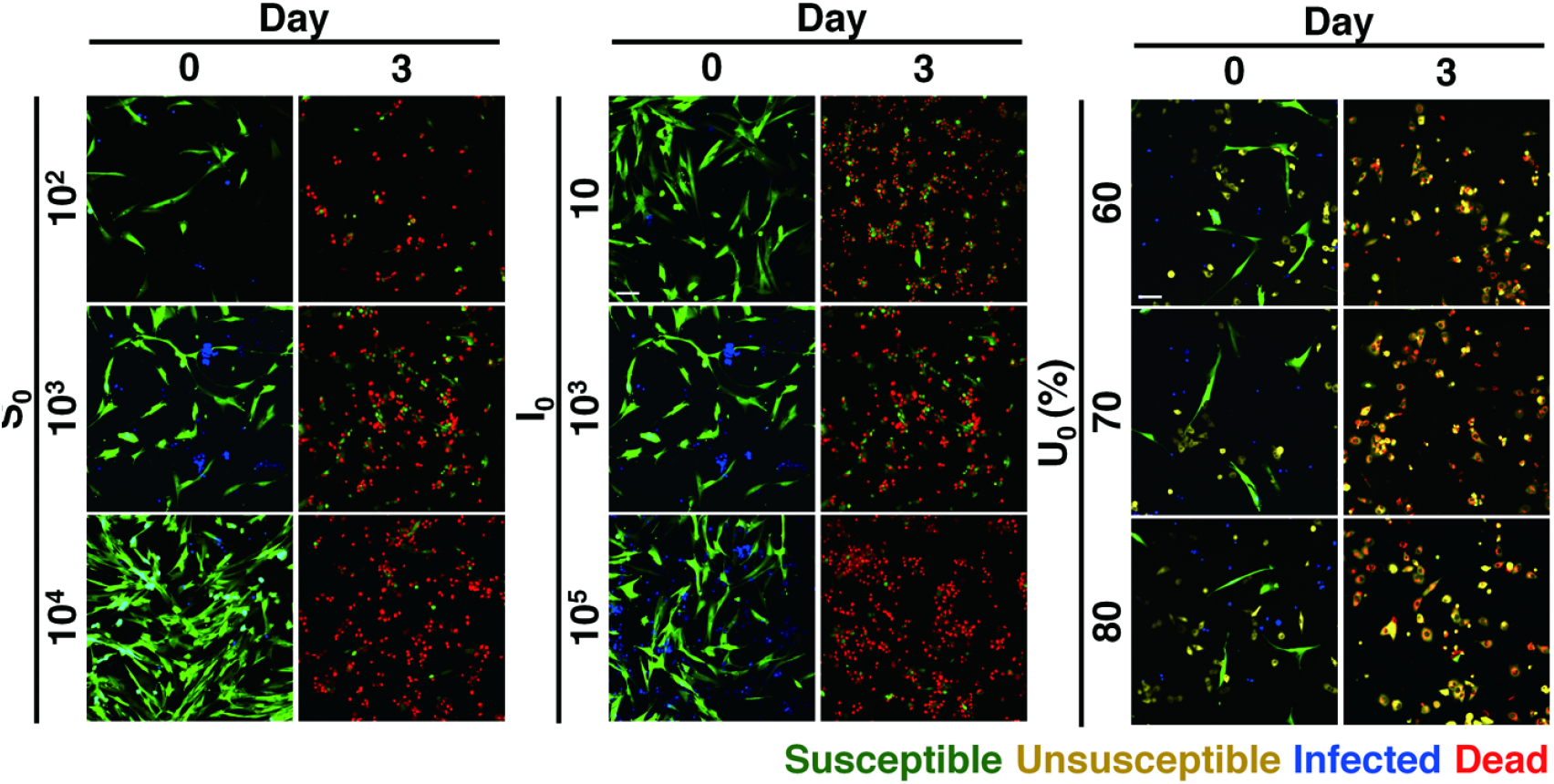
Effects of S_0_ and I_0_ cell densities and U_0_ proportions on virus transmission in a 96-well plate on Day 0 and Day 3. **(A)** Susceptible (S_0_ = 10^2^, 10^3^ and 1 0^4^) at I_0_ = 10^3^. (**B**) Infected (I_0_ = 10, 10^3^ and 1 0^5^) at S_0_ = 10^3^. (**C**) Unsusceptible (U _0_ = 60, 70, and 80%) at uni nf e c t e d (S _0_ +U _0_) = 1 0^3^ a nd I_0_ = 10^3^.

**Fig. S6.**
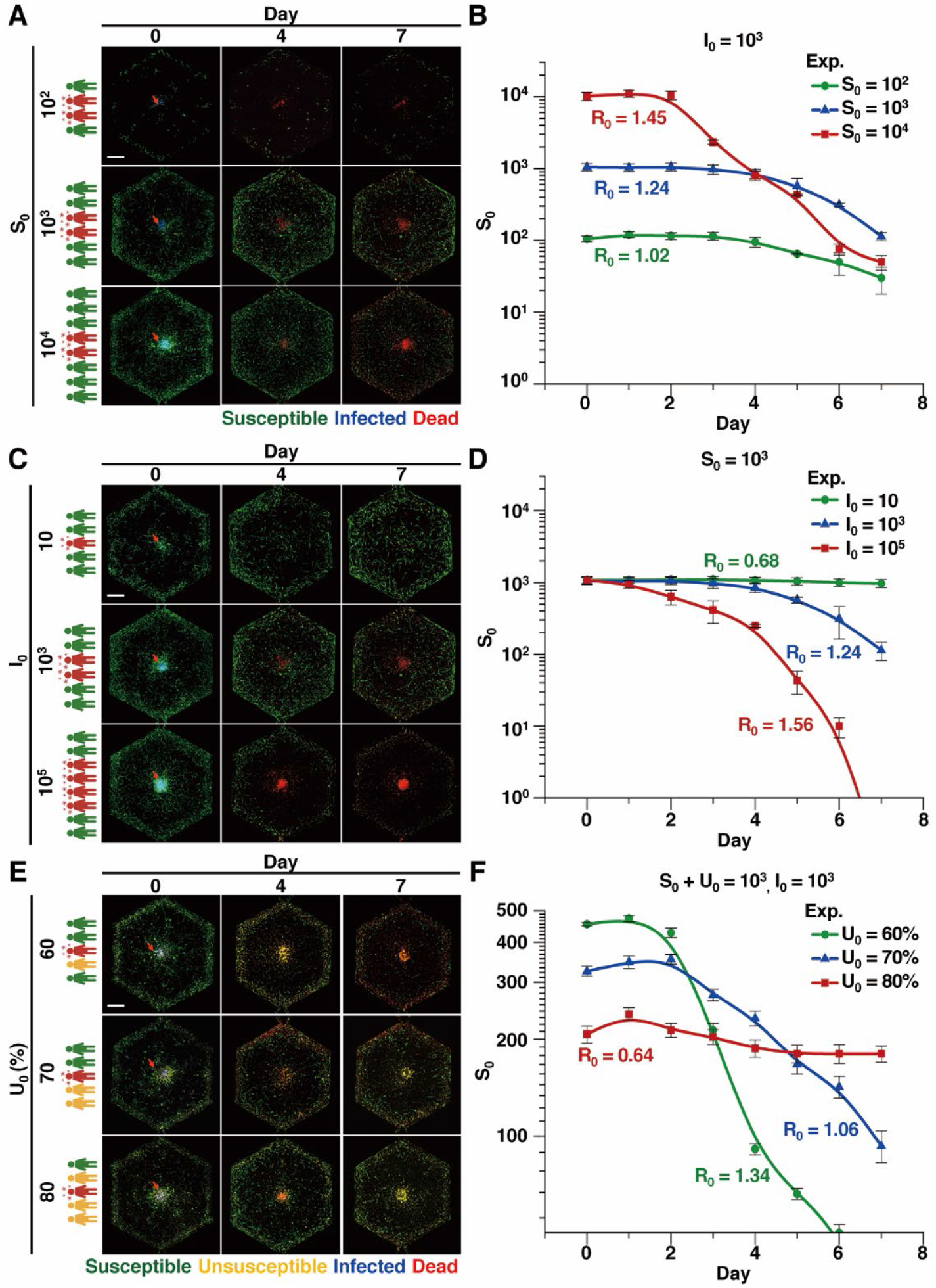
Effects of susceptible, infected, and unsusceptible populations (human hepatocarcinoma Huh-7 cells) on virus (HCoV-229E) transmission. **(A)** S_0_ (10^2^, 10^3^ and 1 0^4^) at I_0_ = 10^3^. **(B)** The S_0_ number (log) for 7 days according to S_0_. **(C)** I_0_ (10, 10^3^ and 10^5^) at S_0_ = 10^3^ on virus transmission. **(D)** The S_0_ number (log) for 7 days according to I_0_. **(E)** U_0_ (60, 70 and 80%) at S _0_ + U _0_ = 10 ^3^ a n d I_0_ = 10^3^. **(F)** The S_0_ number (log) for 7 days according to U_0_. Scale bars = 1 mm. Exp. = experimental results. Sim. = simulation results. Data represent the mean ± S.D.

**Fig. S7.**
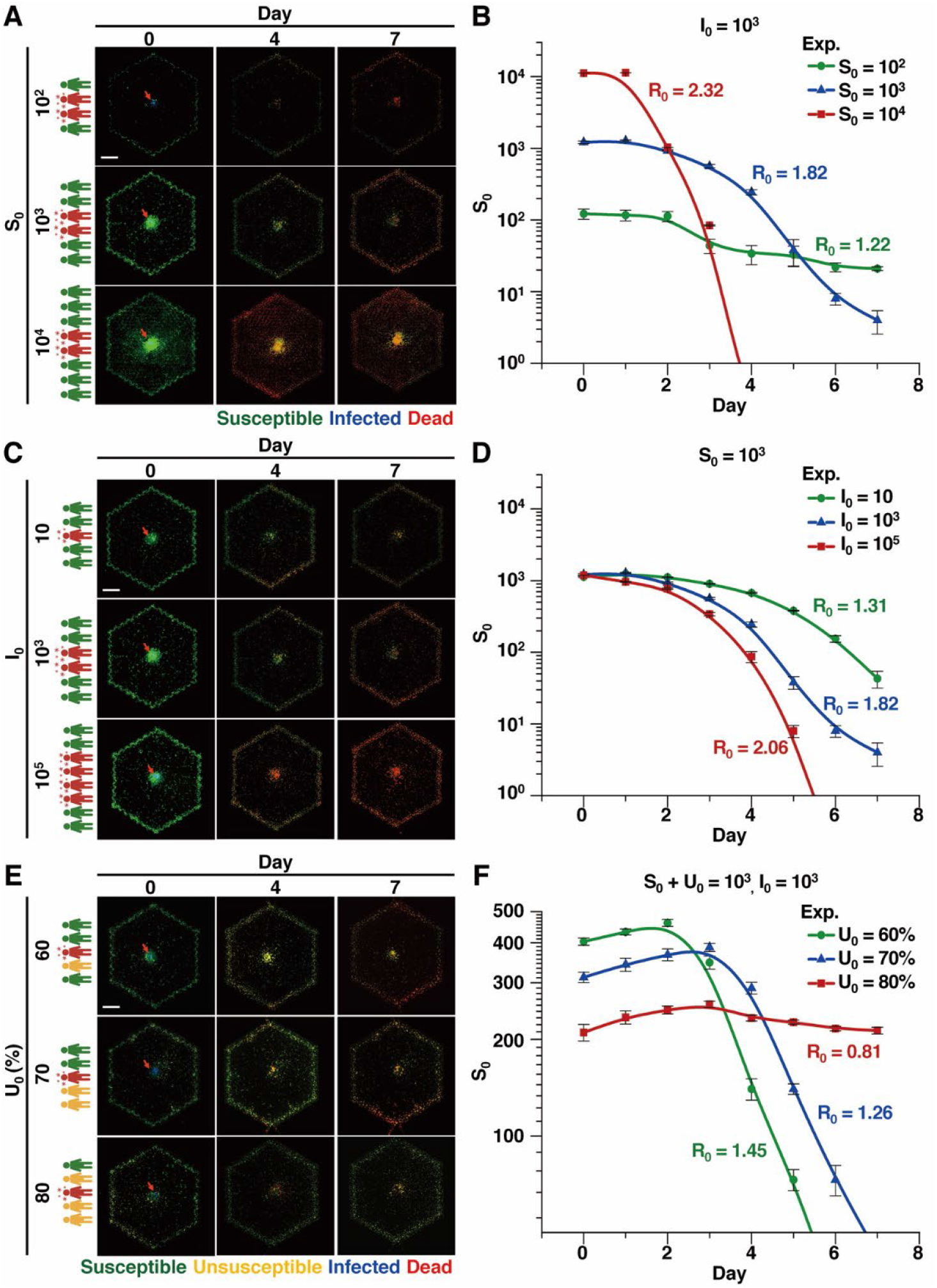
Effects of susceptible, infected, and unsusceptible populations (MRC-5 cells) on virus (HCoV-OC43) transmission. **(A)** S_0_ (10^2^, 10^3^ and 10^4^) at I_0_ = 10^3^. **(B)** The S_0_ number (log) for 7 days according to S_0_. **(C)** I_0_ (10, 10^3^ and 10^5^) at S_0_ = 10^3^. **(D)** The S_0_ number (log) for 7 days according to I_0_. **(E)** U_0_ (60, 70, and 80%) at S_0_ + U_0_ = 10^3^ and I_0_ = 10^3^. **(F)** The S_0_ number (log) for 7 days according to U_0_. Scale bars = 1 mm. Exp. = experimental results. Sim. = simulation results. Data represent the mean ± S.D.

## Notes

### Competing Interest Statement

The authors have declared no competing interest.

